# Pox-AbDab: the Orthopoxvirus Antibody Database

**DOI:** 10.1101/2025.08.22.671799

**Authors:** Henriette L. Capel, Eric Ji Da Wang, Benjamin H. Williams, Charlotte M. Deane, Matthew I. J. Raybould

## Abstract

In August 2024, the World Health Organisation declared the mpox orthopoxvirus to be a Public Health Emergency of International Concern for the second time in three years, emphasising the need for continued studies into its microbiology and potential therapeutic interventions. Here, we present the Orthopoxvirus Antibody Database (Pox-AbDab), a repository of data on antibodies known to bind or neutralise viruses from the same genus as mpox (https://opig.stats.ox.ac.uk/webapps/poxabdab). Beyond standardising and centralising the data, we highlight challenges in translating knowledge across orthopoxviruses, such as the absence of a function-based nomenclature for virion surface antigens. We also performed an exploratory analysis of the known orthopoxvirus-binding antibody landscape, highlighting their aggregate molecular properties, cross-binding/cross-neutralisation profiles, evidence for immunodominance or immune escape from their epitopes, and gaps in coverage to help orient future research.

## Introduction

Mpox is a zoonotic viral disease caused by the monkeypox virus (MPXV), a species of orthopoxvirus first identified in monkey populations in 1958 (1). Mpox infection and transmission in humans has been documented since the 1970s, associated with sudden but relatively localised outbreaks throughout the continent of Africa, containable with standard epidemic response measures (2).

However, a sudden and rapid spread of mpox in May 2022 to all six WHO global regions resulted in the first declaration of an mpox outbreak as a Public Health Emergency of International Concern (PHEIC). In Summer 2024, a genetically divergent mpox upsurge in the Democratic Republic of the Congo spread intercontinentally and led to its second declaration as a PHEIC by the WHO. By 27^th^ July 2025^1^, 48,862 lab confirmed cases have been reported across Africa since 2022, of which 28,953 since 2025. Together, the pattern of more frequent explosive viral spread, PHEIC declarations, and viral genetic divergence stress the ongoing threat of poxviruses and the need for preparedness measures.

During the COVID-19 pandemic, we compiled and released the Coronavirus Antibody Database (CoV-AbDab) containing coupled molecular and phenotypic data on antibodies and nanobodies that can bind to at least one betacoronavirus (3).

Researchers have continually leveraged this centralised information to profile natural antiviral immune responses, assess the impact of vaccination on response shaping, make predictions on potential trajectories of viral escape, and guide the design of monoclonals, bispecifics, or cocktails as prophylactics/therapeutics (e.g. 4–6). An analogous database of antibodies that can bind or neutralise orthopoxviruses would help orient similar investigations from the emergence of any future mpox outbreak. However, while CoV-AbDab provides a helpful template for the design of an antiviral antibody database, substantial adaptations to the data structure are required to account for orthopoxvirus molecular biology.

Orthopoxviruses belong to the genera of poxviruses and contain four species that are pathogenic for humans: variola virus (VARV), vaccinia virus (VACV), cowpox virus (CPXV), and MPXV (7). Unlike single-stranded RNA betacoronaviruses, poxviruses are double-stranded DNA viruses. The central region of the genome is highly conserved and encodes proteins involved in replication and assembly (8), while the flanking regions are variable between virus species and related to host-pathogen interactions (7). For instance, these regions of the left and right termini encode proteins that distinguish MPXV clade I (which caused the 2024 outbreak) from clade II (which caused the 2022 outbreak) (9, 10). Missing and truncated genes between the two clades result in distinct epidemiological features (11, 12); clade I has a higher morbidity/mortality rate and exhibits greater human-to-human transmission compared to clade II (9).

In contrast to betacoronaviruses, which only use a single virion type, the orthopoxviruses have two types of infectious virions which enable replication and maturation in the cytoplasm of host cells: the extracellular envelope virus (EEV) and the intracellular mature virus (MV). The MV is the most abundant and important for inter-host transmission (13). Cell lysis allows MVs to be present outside the cell and infect new hosts by direct fusion with cell membranes (14). Additional packing of intracellular MVs in the cytoplasm results in intracellular envelope virus (IEVs) that are secreted as EEVs through cell-associated envelope virus (CEV) (15). Therefore, EEVs have an additional membrane that needs to be disrupted prior to cell entry, and are important for intra-host transmission (16, 17).

Further complexity is added by MV and EEV bearing distinct sets of surface proteins. More than 20 surface proteins have been identified for orthopoxviruses with the majority located on the MV (18). At least four (A26, A27, D8, and H3) are associated with surface attachment and eleven (A16, A21, A28, F9, G3, G9, H2, J5, L1, L5 and O3) form the entry fusion complex (EFC) (see Table S1).

Of these proteins only a subset – MV proteins A27 (attachment), D8 (attachment), F9 (EFC), H3 (attachment), and L1 (EFC), and EEV proteins A33 and B5 (both involved in viral spread) – are considered immunodominant for the B-cell response (19), although other proteins have been shown to elicit some form of antibody response (20, 21). The complement system (22) is often essential for neutralising activity of these antibodies (23–26), another distinction from anti-coronavirus antibodies which typically neutralise *via* direct inhibition of spike to ACE-2 (3).

Here, we present Pox-AbDab, a freely-accessible repository that centralises sequence, structural, and functional data on antibodies against orthopoxviruses. We show how Pox-AbDab can reveal knowledge gaps and contribute molecular context to debates around the most effective health protection strategies against orthopoxviruses and the pressures driving their antigenic drift.

## Results

### Pox-AbDab stores sequence, structural, and functional data of antibodies against orthopoxviruses

As a first step to building Pox-AbDab, we explored to what extent existing resources contain variable region (Fv) sequences of antibodies that engage orthopoxviruses. We previously built PLAbDab (27) and PLAbDab-nano (28), which respectively scrape all antibody and nanobody sequences deposited in Genbank and provide a tag to all common antigens mentioned in their source literature. By searching for a variety of orthopoxvirus-related keywords (see Methods) and reading the studies to verify specificity to an orthopoxvirus, we were able to retrieve 160 antibodies from PLAbDab. Alternative publicly available database searches revealed 22 relevant entries in the IEDB (29). Combined, this search resulted in 175 unique entries, which we added to Pox-AbDab. To further extend our database, we developed queries to identify orthopoxvirus-related papers (see Methods) and manually mined these to determine whether variable region sequence information is provided.

We were able to retrieve a further 133 entries; as of June 2025, Pox-AbDab contains 308 antibodies or nanobodies (264 and 44, respectively) that bind at least one of the four most studied orthopoxviruses that can infect humans (Figure S1A).

These antibodies are mainly derived from human repertoire B cells (Figure 1A), and focus has recently shifted from VACV to MPXV-specific antibodies (Figure 1C). This aligns with general orthopoxvirus literature following the PHEIC declarations and the long-standing use of VACV as a model virus (30). The data are primarily curated from a relatively small set of sources each reporting a sizeable batch of antibodies or nanobodies (Figure 1C). Full variable region sequences are available for 62% entries (Figure 1B) while 15 antibodies are further characterised with a crystal structure (Figure S1D). Further general statistical breakdowns of the data are available in Figure S1.

**Figure 1.**
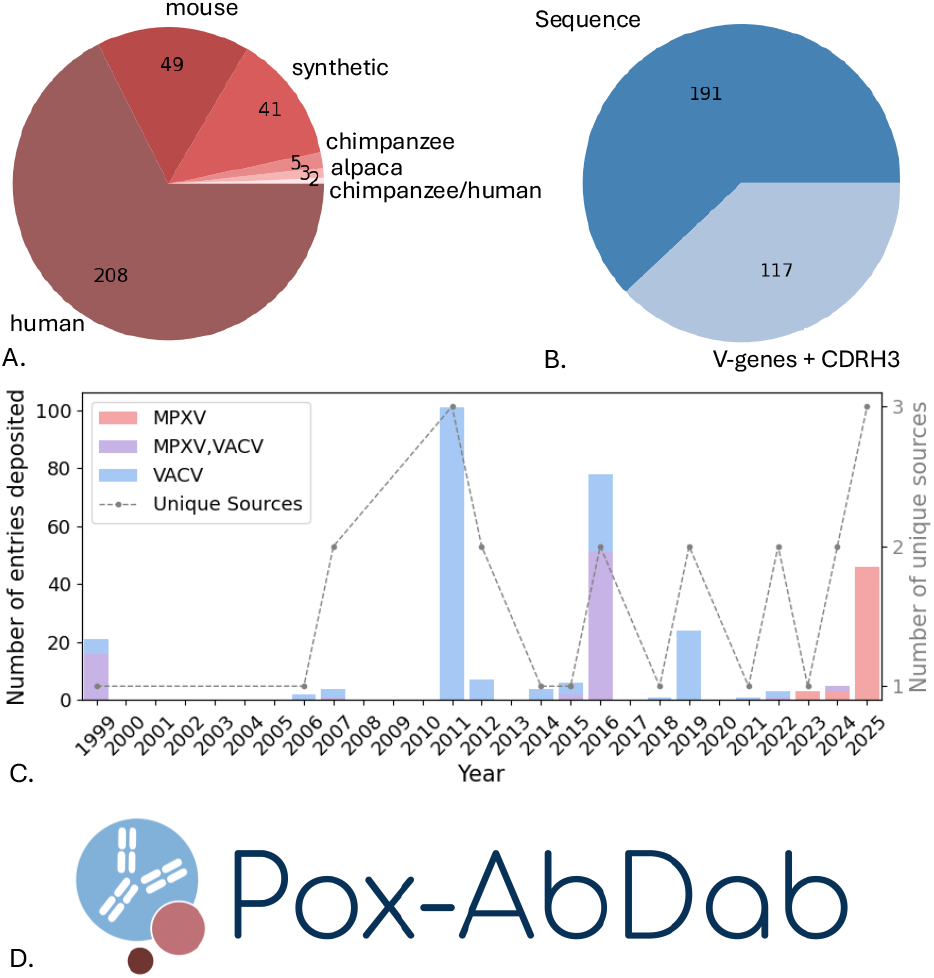
Pox-AbDab database content and Web Application. (A) Species origins of antibodies stored in Pox-AbDab. Most antibodies are derived from humans. (B) Available sequence information of antibodies stored in Pox-AbDab. For 62% of the entries full sequence information is available. (C) Deposited antibodies stored in Pox-AbDab and their sources per year. Model virus VACV is historically most studied, while recent shifts to MPXV are observed. (D) We release Pox-AbDab as a freely available web application (https://opig.stats.ox.ac.uk/webapps/poxabdab).

All the data in Pox-AbDab are available through a web application, hosted at https://opig.stats.ox.ac.uk/webapps/poxabdab. This can be searched by antibody properties (construct, species origin, available epitope information, V- and J-genes), antigen properties (virus species or strain, cellular location), or for particular binding, neutralisation, or protection profiles (see Figure S2). A table with sequence similarity scores for antigen proteins stored in Pox-AbDab is hosted on the main page, to help users find all entries relevant to their research question. The complete database can be downloaded for offline analysis.

### Pox-AbDab antibodies target most immunodominant antigens but gaps exist to functionally-essential antigens

We next investigated the coverage of orthopoxvirus antigens in Pox-AbDab. Orthopovirus antigens are frequently labeled according to a nomenclature inspired by the genome of the VACV Copenhagen strain prior to the era of whole genome sequencing. In this scheme, proteins are annotated with a HindIII restriction endonuclease DNA fragment letter followed by an Open Reading Frame number (7). As it is solely based on genome organisation, this nomenclature can lead to common orthopoxvirus antigens that have the same function being assigned different names. This is even the case among strains of the same virus; for example, orthologs of VACV antigen B5 are labeled B6 in the Bangladesh-1975 strain of VARV and B7 in the India-1967 strain. Some papers replace the virus-specific nomenclature with a label that maps the antigen to an orthologous protein in VACV.

For consistency, we label antigens in Pox-AbDab with the annotation supplied in the source paper. For the purposes of this analysis we have additionally highlighted orthologous proteins across orthopoxviruses by colour (Figure 2). For example, the same colour is used to identify orthologs named after the VACV protein (e.g. VACV A33 and VARV A33(o)) while a slightly lighter shade of the same colour is used when an antigen is given a virus-specific name (e.g. VARV A36).

**Figure 2.**
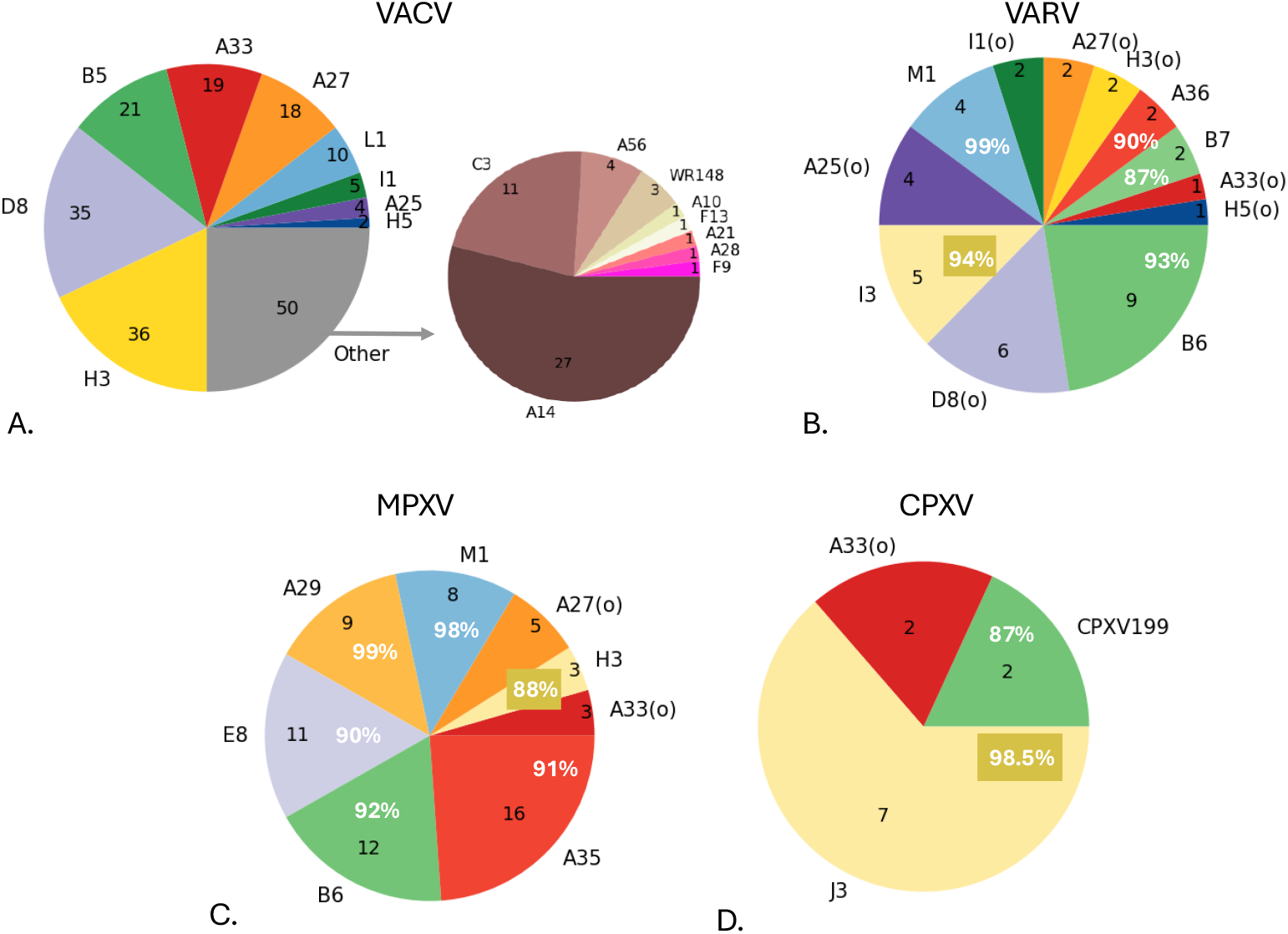
Antigen coverage for antibodies in Pox-AbDab. The number of antibodies against various antigens in VACV (A), VARV (B), MPXV (C), and CPXV (D). Antigens are shown according to their labeling in Pox-AbDab and coloured based on studied homology. Most VACV proteins are labelled according to the Copenhagen strain, while orthologs in the other viruses can be labelled according to the VACV ortholog (indicated with “(o)”) or their own naming scheme. The former is assigned the identical colour used for VACV, the latter is assigned a slightly lighter shade of that colour, and includes the sequence identity score to VACV (in white). The sequence similarity scores were determined based on the Genbank IDs OP868847.1 (VACV), DQ441428.1 and DQ437581.1 (B6-only) (VARV), HM172544.1 (MPXV), and AF482758.2 (CPXV199) and X94355.2 (J3) (CPXV).

Figure 2 shows that Pox-AbDab antibodies target a variety of antigens and, as most entries in the database derive from human B cells, highlights the wide immunogenic response to orthopoxvirus infections or vaccinations. The VACV proteins H3, D8, B5, A33, A27, and L1, which have previously been characterised as immunodominant (19, 31), are correspondingly part of the most abundant proteins in Pox-AbDab. However, antibodies binding other antigens with essential viral functions are scarce or completely lacking (see Table S1).

### Orthopox cross-reactivity is common but not universal across Pox-AbDab antibodies

Orthopoxvirus genomes display over 90% nucleotide identity to one another (7, 32). This is, for example, considerably higher than the similarity between SARS-CoV and SARS-CoV-2 (79.6%) (33) and would imply an even greater potential for antibody cross-reactivity across orthopoxviruses than across betacoronaviruses. Binding and neutralisation assays from the source literature (mainly directed to VACV) show many Pox-AbDab antibodies do indeed exhibit cross-reactivity to several orthopoxviruses (Figure 3A and Figure 3B). However, despite these high levels of homology across orthopoxviruses, antibody cross-reactivity is by no means guaranteed. For instance, Pox-AbDab antibodies binding to VACV are found also to bind MPXV in 77% of the tested cases. For cross-neutralisation, this statistic drops to just 44% (Figure 3C and Figure 3D), which may suggest that mpox-specific vaccination approaches offer a benefit over the current vaccinia-based vaccinations suggested to protect against MPXV, in terms of inducing robust mpox immunity. An activated complement system was required for effective neutralisation by at least 45 antibodies in Pox-AbDab (Figure S1F); the requirement for efficient recruitment of other components of the immune system may add complexity to neutralisation potential beyond the capacity to cross-bind.

**Figure 3.**
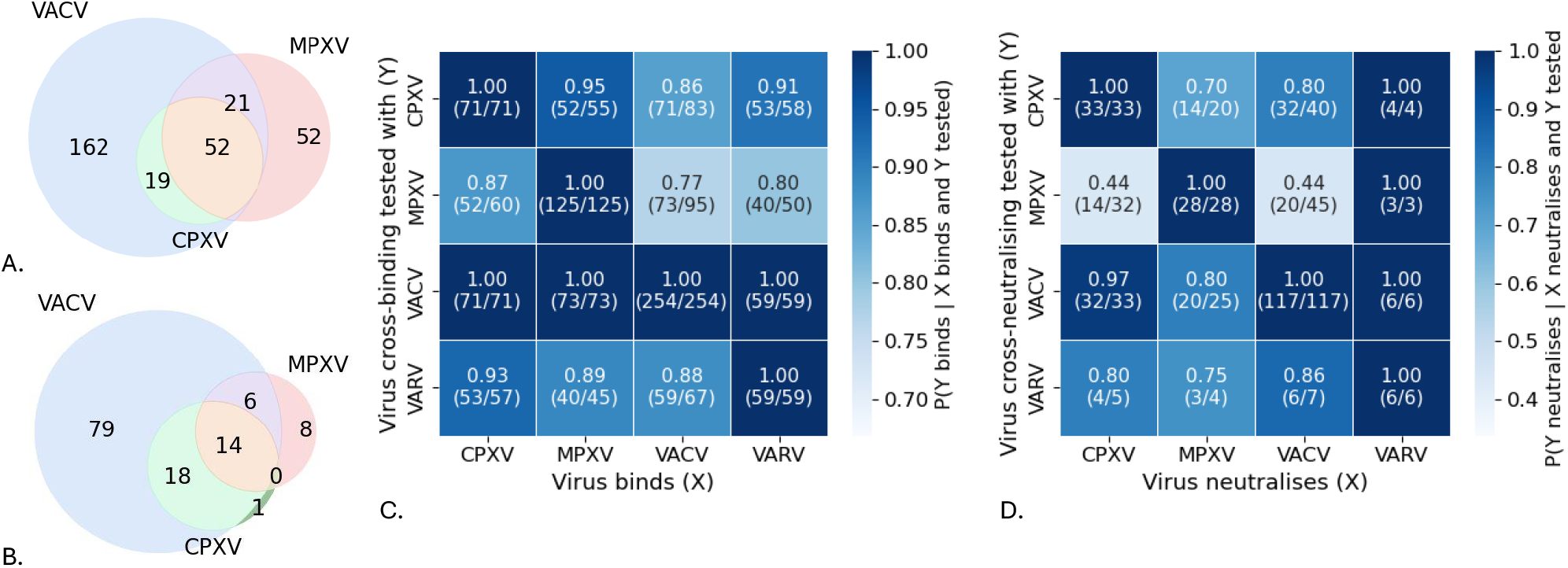
Cross-binding and cross-reactivity between orthopoxviruses. The number of cross-binding (A) and cross-neutralisation (B) antibodies in Pox-AbDab against VACV, MPXV, and CPXV. Fraction of cross-binding (C) and cross-neutralisation (D) of antibodies in Pox-AbDab. The number of antibodies considered are shown in brackets. While all antibodies binding to VARV, MPXV, and CPXV tested against VACV bind VACV as well, not all VACV targetting antibodies cross-bind to the other viruses, and cross-neutralisation is even rarer.

### Structural data in Pox-AbDab reveals the potential for immune evasion

Antibodies can impart a natural selection pressure on viruses to mutate so as to evade the protective immune response of their hosts. Such immune evasion is particularly evident in betacoronaviruses, where even early SARS-CoV-2 strains contained mutations that interrupted binding to common human antibody epitope clusters (34). Orthopoxvirus antigens generally show high sequence similarity between species and strains, but it remains unclear whether the variation that does exist is driven by immune escape (e.g. from the vaccinia vaccine), driven by another pressure, or simply reflects genetic drift.

We analysed the binding footprints of the human antibodies structurally characterised in Pox-AbDab, exploring how conserved these binding residues are across orthopoxviruses (Table 1). This revealed a mixed picture dependent on the surface antigen targeted. The epitopes of seven human antibodies against A27, L1, and M1 were entirely conserved across the orthopoxvirus orthologs, even in binding footprints spanning 22 residues. However, eight human antibodies against A33, B6, and D8 targeted sites that vary in sequence across the examined orthopoxviruses. For instance, the D8 epitope footprint for the antibody in 5USH is only 75% conserved. Some positions differ markedly in amino acid chemistry (e.g. the A33 epitope residue 168 in crystal structure 4M1G is negatively charged in VACV, CPXV, and VARV, but positively charged in MPXV).

**Table 1.**
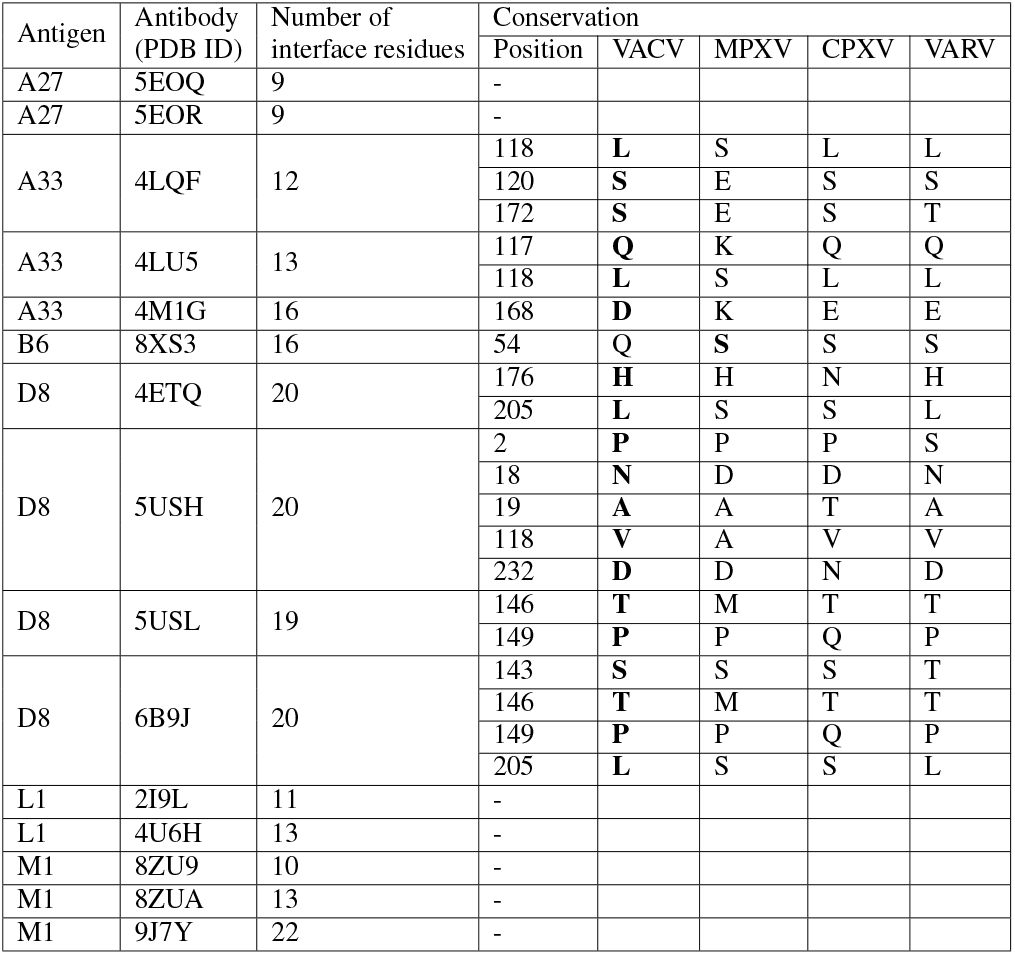
Epitopes for the 15 antibodies in Pox-AbDab with solved crystal structures. Any non-conserved residues in the binding footprint across the four orthopoxvirus species are shown; virus sequence definitions: VACV (OP868847.1), MPXV (HM172544.1), CPXV (AF482758.2), and VARV (DQ441428.1). The residue in the epitope of the solved crystal structure is shown in bold. Antigens are labeled according to the VACV Copenhagen nomenclature.

As a further case study, we analysed Borealpox (BRPV), the most recent orthopoxvirus known to infect humans (35). Borealpox is not yet included in Pox-AbDab as currently no sequences of antibodies against BRPV have been reported in the literature. Here, we consider its B6 ortholog. Borealpox B6(o) displays relatively low sequence identity to VACV B5 (58%) and yet despite this divergence, its AlphaFold3 (36) model retains the same overall fold as the crystallized fragment of MPXV B6 and VACV B5, suggesting functional conservation (Figure 4A). We then considered where the non-conserved residues lie in relation to the antigen-binding site of antibody hMB668 (37), an mpox-reactive clone derived from a vaccinia-vaccined individual (Figure 4B). Across the 15-residue epitope, just 5 residues are conserved between MPXV and BRPV, or between VACV and BRPV (Figure 4C).

**Figure 4.**
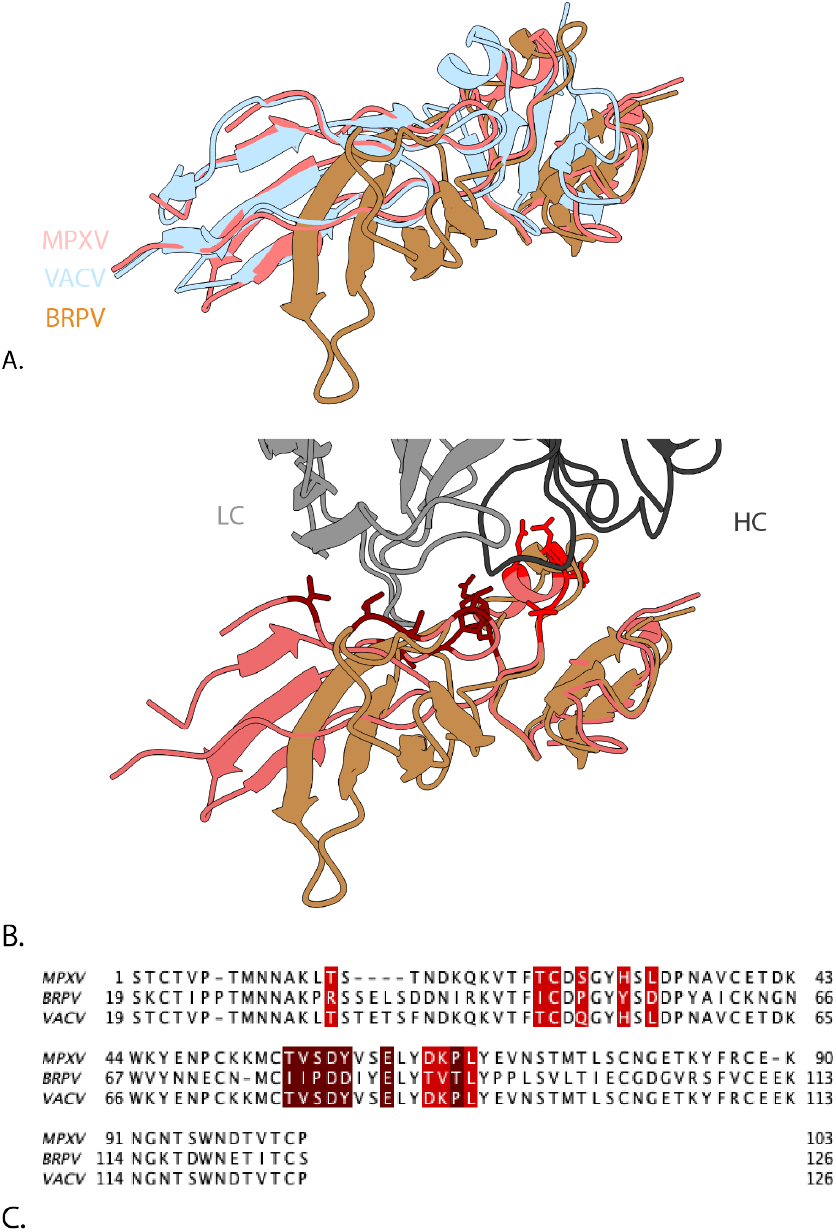
The crystal structure of antibody hMB668 (named D68 in the PDB) bound to mpox B6(o) (PDB ID: 8XS3). (A) The MPXV B6 protein from 8XS3 (light red) aligned to AlphaFold3 predictions of VACV B5 (light blue) and the BRPX ortholog (orange). (B) Residues on MPXV B6 interacting with antibody hMB668 (grey) shown in dark red (heavy chain) and red (light chain). (C) Sequence alignment between MPXV B6, BRPV B6(o) and VACV B5 with the interacting residues highlighted (heavy chain footprint in dark red, light chain footprint in red).

### Genetic profiling of human Pox-AbDab antibodies currently offers little evidence for which clones are immunodominant

Finally, we assessed the sequence diversity and antigen-binding convergence of antibodies in Pox-AbDab. Antibodies in Pox-AbDAb span a full range of germline and CDR3 sequence length usages as observed in natural sequences (see Figure S3, Figure S4, and Figure S5). The germline-likeliness of these antibodies was explored by comparing the 130 human antibody sequences in Pox-AbDab against a set of natural human naive B cell receptor repertoires from the Observed Antibody Space (OAS) (38, 39) database using the KA-Search tool (40) (Methods, Figure 5), looking for the most sequence identical match. This revealed that the human orthopoxvirus antibodies in Pox-AbDab antibodies on average lie some distance away from the naive baseline repertoire. As a reference, we also assessed the proximity of a random sample of 130 human B-cell-derived coronavirus-binding antibodies from CoV-AbDab (3) (Figure 5). While the Pox-AbDab clones had median closest chain identity of 86% (heavy chain) and 94% (light chain), the CoV-AbDab clones sat closer to the naive repertoire with 92% (heavy chain) and 97% (light chain). We obtained comparable results when comparing to all B cell receptors in OAS (Figure S6).

**Figure 5.**
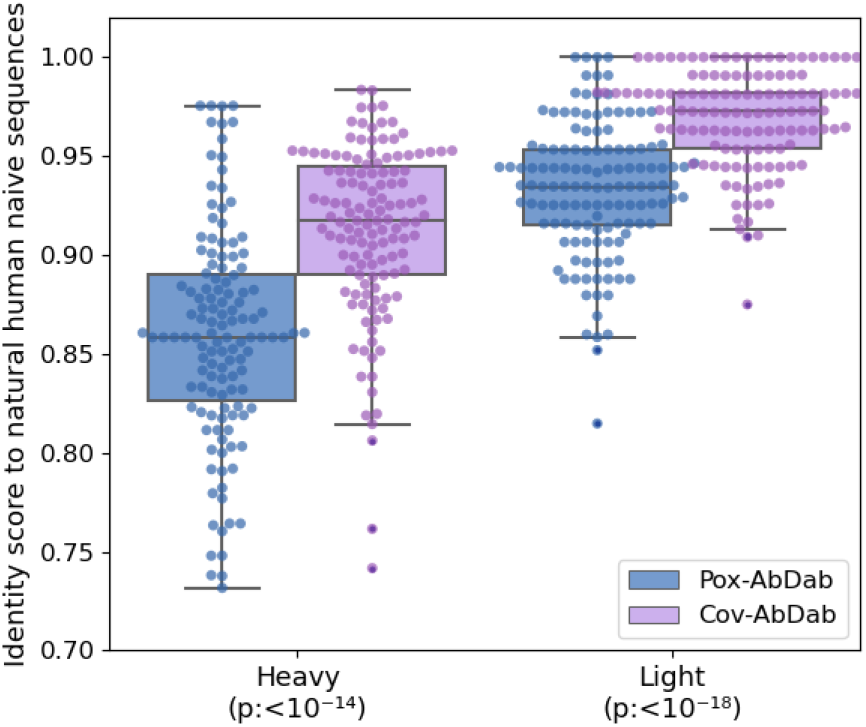
Antibody heavy and light sequence identity score to a set of human antibodies from naive B-cells. The 130 full-length human antibody sequences in Pox-AbDab (blue) and 130 randomly selected human antibody sequences from CoV-AbDab (3) (purple) were compared against human naive B cell receptor sequences from OAS (38, 39) (28.3M heavy chains from 12 studies, 24.6M light chains from 5 studies). The highest sequence identity observed was recorded. P-values of the Mann–Whitney U test are indicated on the x-axis.

We surveyed the Pox-AbDAb clones for evidence of response convergence. We first applied clonotyping, which groups antibodies by usage of the same heavy chain V/J genes and that bear length-matched and highly similar CDRH3 loop sequences. We obtained multi-occupancy clusters that comprised a total of 62 antibodies, however all antibodies within the same cluster were derived from the same source, and though the libraries contained pooled blood from many individuals these clones could derive from the same donor.

It has been frequently observed that antigen-specific antibodies with similar Fv loop shapes tend to engage the same epitope (41). We therefore applied SPACE2, a method for 3D modeling and clustering of antibodies with high Fv structural homology. The 147 antibodies in Pox-AbDab with complete VH and VL sequences clustered into 16 multi-occupancy clusters (119 clusters in total); when allowing clustering between antibodies with length-dissimilar CDRs, this grew to 19 multi-occupancy clusters. Again, all clusters contained only antibodies reported from the same source. Overall, this indicates that Pox-AbDab cannot yet confidently tell us which clones are immunodominant upon orthopoxvirus infection or vaccination.

## Discussion

Pox-AbDab currently stores 308 antibodies binding to at least one of the orthopox virus species VACV, VARV, MPXV, and CPXV. For context, this represents roughly three times more data than could be scraped from the literature on betacoronavirus binders prior to the SARS-CoV-2 pandemic (3). New data are continually emerging (e.g. Yefet *et al*. (42), whose sequence and structural data should be released post peer review) and we will continuously capture these through routine maintenance.

Poxvirus binding and heavy chain clone information are available for every entry, and where possible we have added neutralisation and *in vivo* protection data, the full Fv region sequence and structural data. Literature sources and notes tagged to each entry provide ready access to additional information on characteristics.

Around 52% of the entries in Pox-AbDab were contained in GenBank and therefore retrieved by PLAbDab (27), showing the utility of this resource in generating initial datasets of target-associated sequences whose antigen specificity can be confirmed by inspection of the source literature. However, we could find substantially more data through manual curation, highlighting the diverse documentation of antibody sequence information across studies, including through supplementary tables and files, and even figures displaying sequence alignments. The sequences of antibodies mentioned in several sources were not obtainable — older papers do not share sequence data as reliably as newer ones, while the sequences of some antibodies commonly used as baselines still have restricted access (e.g. Mab 69-126-3-7 and 1G10), or are restricted from reusage/distribution (43).

A particular challenge underpinning the need for manual curation was the heterogeneity in both virion and antigen naming, as independent studies frequently refer to the same viral protein by its VACV ortholog and virus-specific nomenclature. We provide a reference table of orthologous protein names alongside the web application to assist new entrants to the field, although a longer term solution would be a community effort to establish a molecular function-based antigen naming scheme. The diversity of species and strain usage for binding, neutralisation, and protection assays, even within a study (44), also pose a challenge for any attempt to extract accurate functional labels in an automated fashion.

As with all literature-based curations of antigen-specific data, biases exist in Pox-AbDab that should be taken into account when interpreting analyses. For example, VACV is the most studied orthopox species and is therefore the most represented virus species among the data in Pox-AbDab. In MPXV-focused studies, neutralisation and protective capacity of the antibody is often tested only in VACV due to data availability and safety regulations around BSL-3 pathogens (37, 45, 46). Biases towards the MV compared to the EEV are caused by the fragile outer membrane of the envelope virion, which is hard to study (18). Other knowledge gaps identified in Pox-AbDab (see Figure 2) do not have obvious origins in regulatory or technical challenges, and might be readily remedied by focused research efforts.

The aggregate data in Pox-AbDab already reveals immunologically relevant patterns amongst human antibodies to orthopoxviruses, such as their tendancy to accrue higher levels of somatic hypermutation than their coronavirus-binding counterparts. We also highlight evidence of sequence variability in human antibody epitopes for certain surface antigens and that only a moderate number of antibodies in the database have been shown to cross-neutralise both VACV and MPXV. As we accrue more orthopoxirus-specific lineages from post-vaccination/post-infection B cell receptor repertoires, immunodominant clones may emerge as they did in CoV-AbDab (3). The presence or absence of such signals would add useful context to our understanding of the potential for orthopoxvirus immune evasion.

As a pandemic preparedness measure, Pox-AbDAb provides an immediate library of molecular designs that could be tested as laboratory reagents or diagnostic, prophylactic, or therapeutic antibodies in the context of an emergent orthopoxvirus outbreak. It also acts as a central location where specific antibody data could be deposited as studies into the immune response to existing or new orthopoxviruses emerge, akin to CoV-AbDab (3). This not only accelerates analyses into the potentially protective properties of these clones, but also provides a ready source of machine learning-grade data that could be leveraged to design new molecules with desired orthopoxvirus reactivity and neutralisation profiles.

## Methods

### Data curation

Academic papers and patents containing antibodies binding orthopoxvirus pathogenic for humans were sourced by querying PubMed, Elsevier Mpox Information Centre, Google Patents, and Google Scholar, and the publicly available maintained databases PLAbDab (47), PLAbDabnano (28), IEDB (29), and INDI (48) with the search terms: orthopox, vaccinia, VACV, variola, VARV, monkeypox, mpox, MPXV, cowpox, and CPXV. Antibodies included in Pox-AbDab bind at least one of these four species pathogenic to humans and at least the antibody V-gene(s) and CDRH3 is available. Various binding assays such as surface plasmon resonance (SPR), bio-layer interferometry (BLI), enzume-linked immunosorbent assay (ELISA), protein microarray, and western blot were considered evidence of binding to capture the largest possible collection of relevant antibodies.

### Antibody identification in Pox-AbDab

Curated antibodies were named according to the source naming convention. In some cases, the last name of the first author in which the antibody was initially described is used as suffix to reduce the risk of duplicate names. In some cases, antibodies with the same variable region sequence are given different names depending on the source; we have labeled these entries with both names. The originating species and the antibody construct were extracted from the paper.

### Antibody functionality captured by Pox-AbDab

To consistently store detailed binding data, virus species, strain, location, and protein information was combined to a single identifier separated with underscores (e.g. VACV_WR_MV_H3) For MPXV, clade information was specified when known (e.g. MPXV-IIb). Common abbreviations were used for concise strain identification. The database stores antibodies against the VACV strains Western Reserve (WR), Lister, Elstree, Copenhagen, Connaught, New York City Board of Health (NYCBOH), Munich 1 (Munich1), International Health Division J (IHD-J), Tiantan, recombinant Tiantan strain expressing GFP (TT-FGP), and the smallpox vaccine strain ACAM2000. MPXV strains Zaire 1979 (Zaire79), Zaire96-I-16, Zaire599, and China-C-Tan-CQ01, VARV strains Solaiman, Bangladesh-1975, India-1967, and India-1967-Ind3a, and CPXV strains Grishak and Brigthon Red (BR) are included in the database. The location placeholder is used for the MV virion, the EEV virion, and to indicate secreted proteins. Proteins are identified according to the antibody paper sourced. VACV protein orthologs are indicated by the suffix “(o)”. For example, MPXV protein A35 is identified as the ortholog of VACV A33 and can be indicated by “A33(o)” or “A35” dependent on the nomenclature in the source. Recombinant proteins are indicated by the suffix “(r)” and translated proteins by suffix “(t)”, when information is provided in the source.

As neutralisation assays are not performed on specific proteins, we used the virus species, strain, and location (again concatenated with underscores) as identifiers. Protection was indicated by virus species and strain. For consistent binding, neutralisation, and protection identifiers question marks (“?”) are placeholders for missing data.

### Antibody sequence information

V-gene and CDRH3 information are the minimal criteria for including antibodies in Pox-AbDab. Where available, VH or VHH and VL sequence information is stored, as well as V-gene, J-gene, and both CDRH3 and CDRL3 sequences. ANARCI (49) was used to identify CDR3 sequences and gene information in case sequence, but no or incomplete gene and CDR information, was available. We excluded allele information for consistency. CDRs are defined based on the IMGT (and Kabat) definition. Complete entries (e.g. JF11 and JE10 (50)) or part of the sequence information (e.g. full VH and VL sequence of A26C7 (19)) was excluded when ANARCI was unable to number the sequence and/or ABodyBuilder2 (ABB2) (51) was unable to structurally model the antibody. Extra sequence information beyond the VH and VL, as determined by ANARCI, was removed. Modified or inconsistent sequences of a single entry in multiple sources were stored as separate entries (e.g. VACV-138-v1 and VACV-138-v2).

### Structural Data

Where available, structure information was stored using the PDB ID (52). When unavailable, ABody-Builder2 structural models are provided. Epitope information was labeled as either conformational or linear based on whether or not epitope information was provided as a contiguous string of amino acids (*i*.*e*. a binding domain or loop) or as a set of sequentially discontinuous residues.

### Web application

The data are stored in a relational database, using MariaDB^2^, with a schema defined using the SQLModel library for Python^3^. The web application communicates with the database by means of a REST API conforming to version 3.1.0 of the OpenAPI Specification^4^. Both the API and the web application itself were created using the FastAPI web framework^5^.

### Database analysis

We summarised the database on current information storage and performed immunoinformatic analysis of the data. The origin and format of the antibody were summarised, as well as the source information. The type of sequence data available, the targeted virus species and antigens, the epitope identified, the ability of cross-neutralisation, and the importance of the complement system were also summarised.

The sequence diversity of the database was analysed based on V-gene usage and CDR length of the antibody per originating species, as well as the targeted proteins. Diversity of the human sequences were compared against a randomly sampled set of 5000 antibodies collected from healthy individuals by Jaffe et al. (2022) (53) as stored in OAS (38, 39).

For structural analysis, the diversity of the targeted epitope within the four orthopoxvirus species was determined. The native epitope was identified in the available PDB structures based on a 1.4 probe radius and a 1.5 BSA cutoff. The epitope footprint as indicated in the solved crystal structure was compared to the ortholog antigens in the other species. Here we used accession numbers OP868847.1 (VACV), DQ441428.1 (VARV), HM172544.1 (MPXV), and AF482758.2 (CPXV).

As a case-study, the sequence-structural conservation of the recently identified Borealpox virus B6 ortholog protein was studied. The crystal structure of MPXV B6 (PDB ID: 8XS3 (37)) was compared against an AlphaFold3 (36) model of the Borealpox B6 ortholog (GenBank entry: MN240300.1). Sequence alignments (by Clustal Omega (54)) and visualisation of sequence conservation were performed using ChimeraX (v 1.10) (55). Interacting residues were defined as residues with a buried surface area *≥* 15Å^2^.

### Immunoinformatics

We further analysed antibody sequence diversity and clustered antibodies using clonotyping, paratyping, and SPACE2 (56) to check for antigen-binding convergence of antibodies in Pox-AbDab.

The 130 available human VH and VL sequences were compared against a naive set of human antibodies using KA-Search (40). The naive dataset contained 5000 randomly sampled unpaired heavy and light sequence from the Observed Antibody Space (OAS) database (38, 39), collected on 21th January 2024. Distances to the full OAS naive and memory dataset were determined based on the provided “OAS-aligned” dataset of KA-Search. To compare distance scores against coronavirus-binding antibodies, 130 human full-length antibody sequences from Cov-AbDab were randomly selected. A non-parametric Mann–Whitney U test was used to statistically compare identity scores observed for Pox-AbDab against CoV-Abdab (3).

Clonotyping was performed on all entries in Pox-AbDab and carried out based on matching IGHV and IGHJ identities, identical CDRH3 lengths and *≥* 80% CDRH3 sequence identitity.

Functional convergence beyond sequence similarity was studied by clustering antibodies on their CDR conformations using SPACE2 (56). The 147 antibodies in Pox-AbDab with a complete Fv sequence were modeled using ABodyBuilder2 (51) and clustered according to the author’s recommendation of 1.25Å either in length-restricted mode (all antibodies in a cluster have identical CDR lengths) or length-unrestricted mode (RMSDs calculated as dynamic time warping distances).

## Data availability

The database is accessible without restriction or registration at https://opig.stats.ox.ac.uk/webapps/poxabdab, and all data within Pox-AbDab can be freely downloaded.

## AUTHOR CONTRIBUTIONS

CMD and MIJR conceptualised the study. HLC, EJDW, and MIJR designed the methodology. HLC, EJDW, and BHW wrote software. All authors analysed the data. HLC and EJDW visualised the data. HLC and MIJR contributed to data curation. HLC wrote the original draft with input from all authors. CMD and MIJR reviewed and edited the manuscript. MIJR supervised the work. CMD provided funding and resources for the project.

## ACKNOWLEDGEMENTS

We acknowledge Gemma L. Gordon for supporting the initial literature curation. We acknowledge helpful conversations with the Immunoinformatics team in OPIG that improved the quality of the work.

---

**Figure S1.**
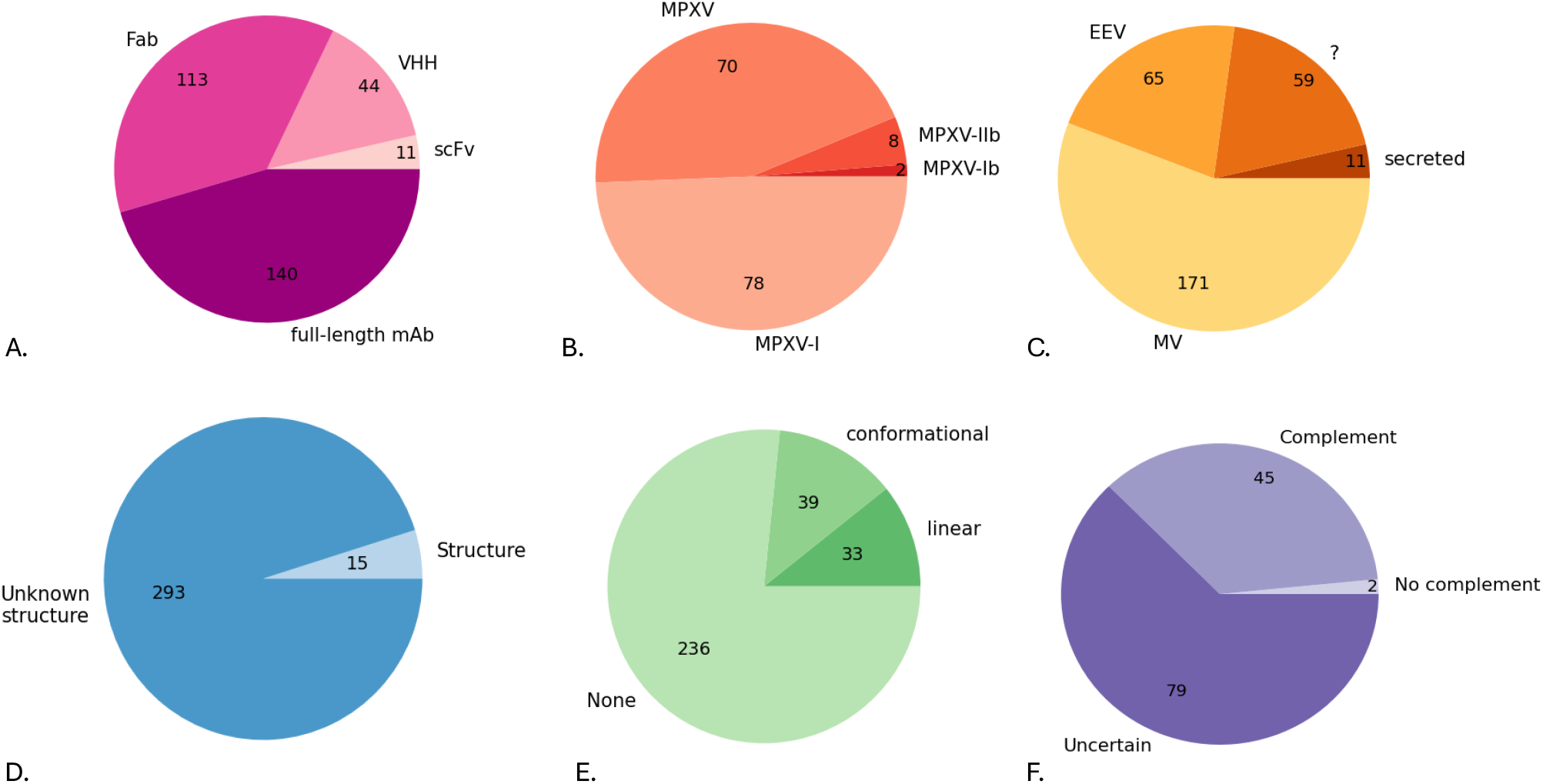
Pie charts showing (A) the ratios of conventional antibodies and single-domain antibodies in Pox-AbDab, with conventional antibodies split by format; (B) where a Pox-AbDab entry has been tested against MPXV, the clade that was tested (if specified, else ‘MPXV’); (C) the location context of each entry, (if specified, else ‘?’); (D) the number of entries for which crystal structures are available; (E) the epitope information available for each Pox-AbDab entry. Linear epitope information is defined as a contiguous string of amino acids. (F) the contingency of neutralisation on an active complement system.

**Figure S2.**
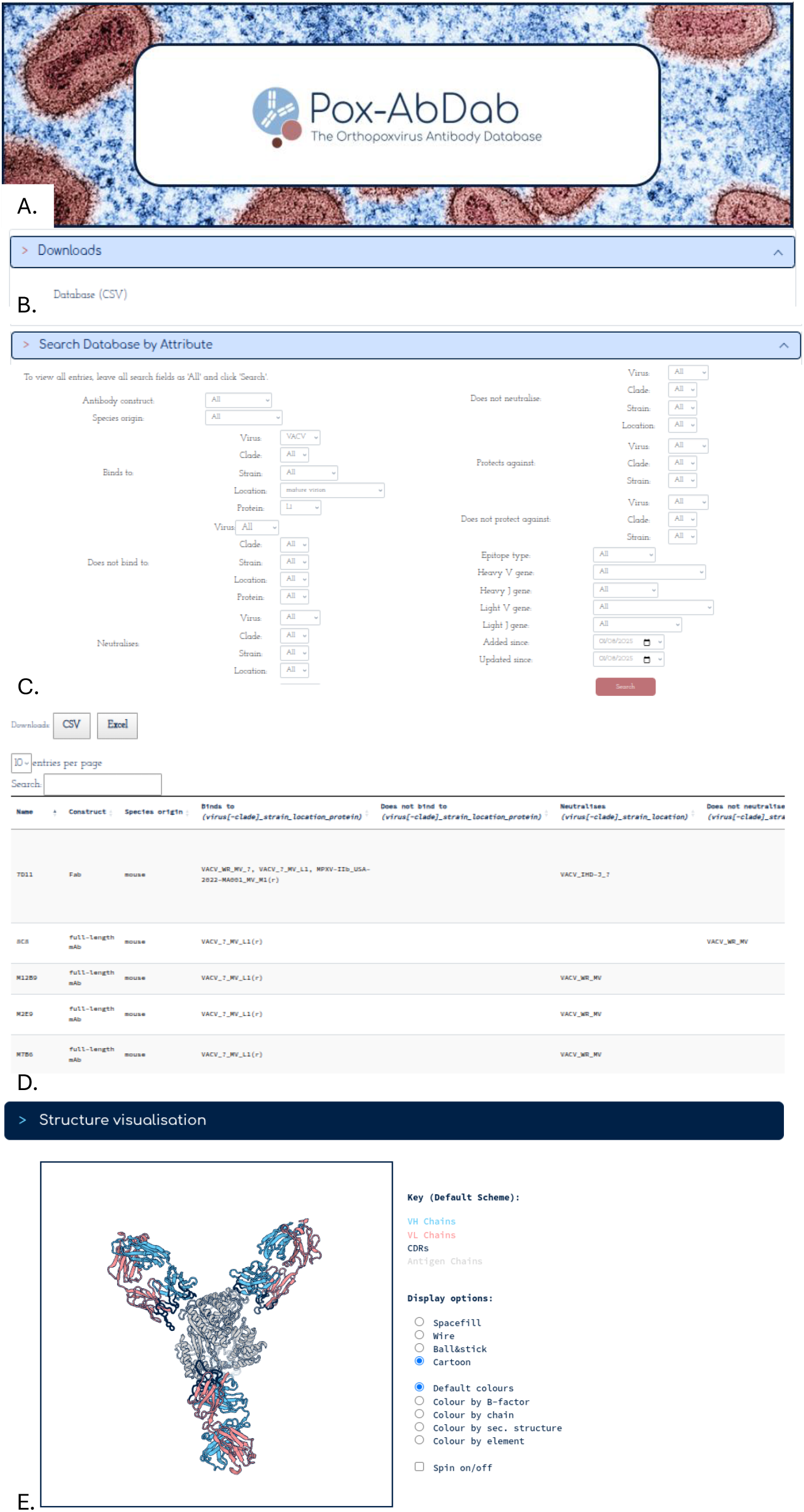
The Pox-AbDab Web Application. (A) The Pox-AbDab homepage logo (background image credit: NIAID, Mpox Virus, CC BY 2.0, https://www.flickr.com/photos/niaid/52988422372/in/photostream/). (B) All Pox-AbDab data can be downloaded. (C) The database can be queried by attribute (neutralisation profile, construct, germlines, etc.). (D) The result table of the attribute search. (E) The result table links the experimental solved antibody structures stored in SAbDab (57) (PDB ID 2i9l visualised here) or modelled structures by ABodyBuilder2 (51).

**Figure S3.**
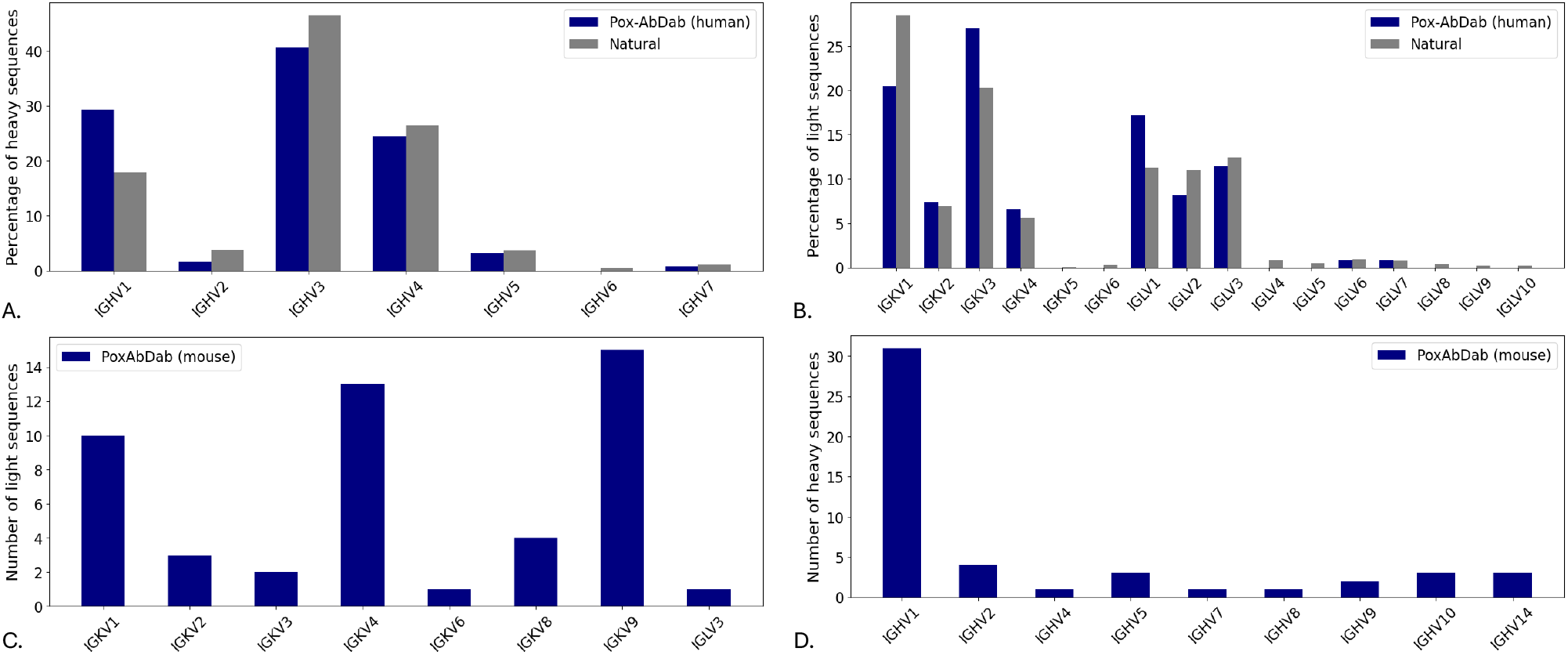
Bar charts showing (A) the distribution of heavy V gene family usage across the human antibodies in Pox-AbDab relative to a sample of the baseline natural repertoire from the Observed Antibody Space (OAS) database (38, 39); (B) the distribution of light V gene family usage across the human antibodies in Pox-AbDab relative to a sample of the baseline natural repertoire from the Observed Antibody Space (OAS) database (38, 39); (C) the distribution of heavy V gene family usage across the murine antibodies in Pox-AbDab; (D) the distribution of light V gene family usage across the murine antibodies in Pox-AbDab. Comparison against the natural human repertoire is based on 5000 randomly samples natural antibody sequence from the Jaffe et al., 2022 (53) as stored in OAS.

**Figure S4.**
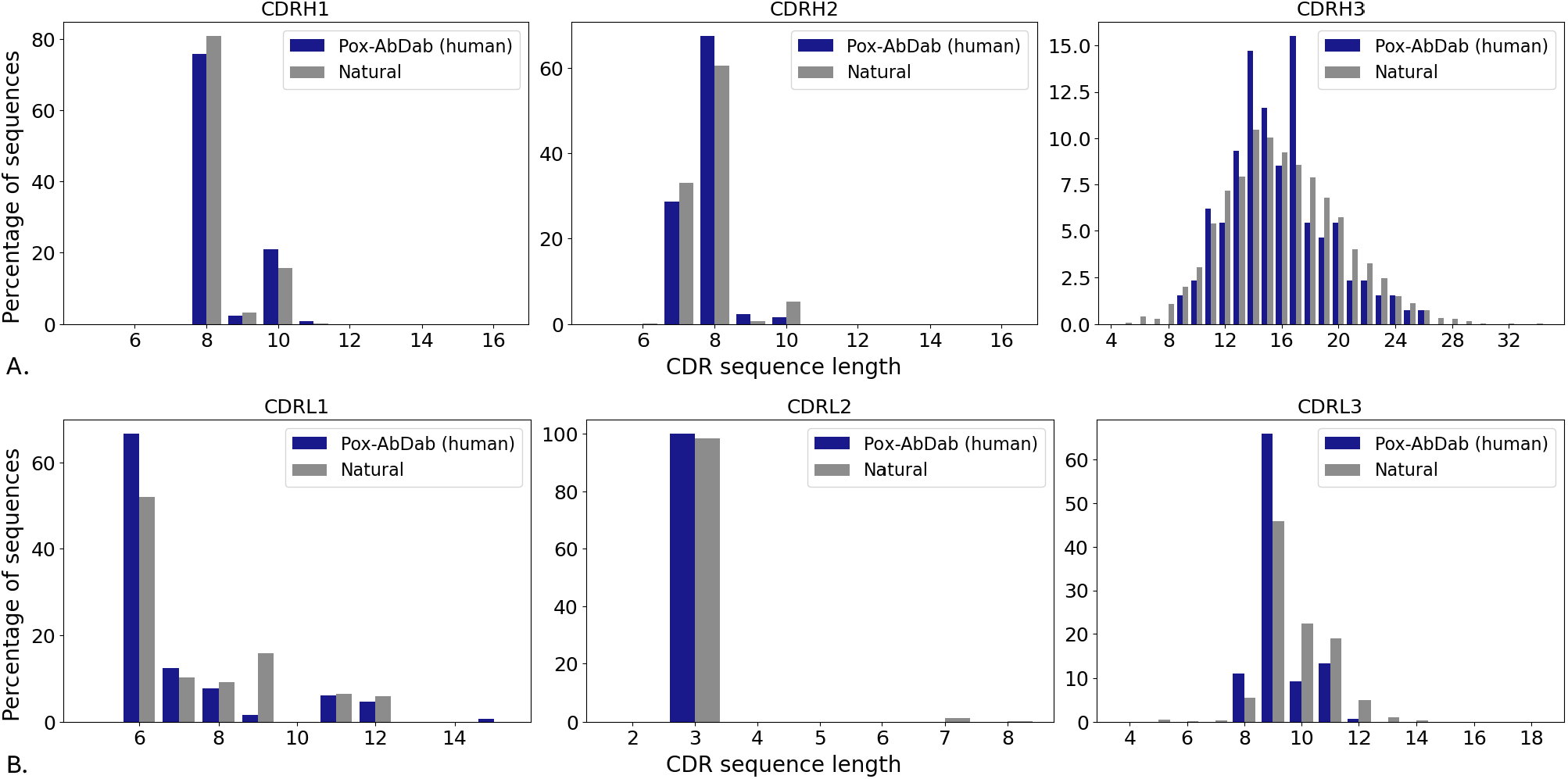
Bar charts showing the distributions of CDR lengths in the full-length sequenced, human antibodies in Pox-AbDab relative to a sample of the baseline natural repertoire from the Observed Antibody Space (OAS) database (38, 39): (A) CDRH1; (B) CDRH2; (C) CDRH3; (D) CDRL1; (E) CDRL2; (F) CDRL3. Comparison against the natural human repertoire is based on 5000 randomly samples natural antibody sequence from the Jaffe et al., 2022 (53) as stored in OAS.

**Figure S5.**
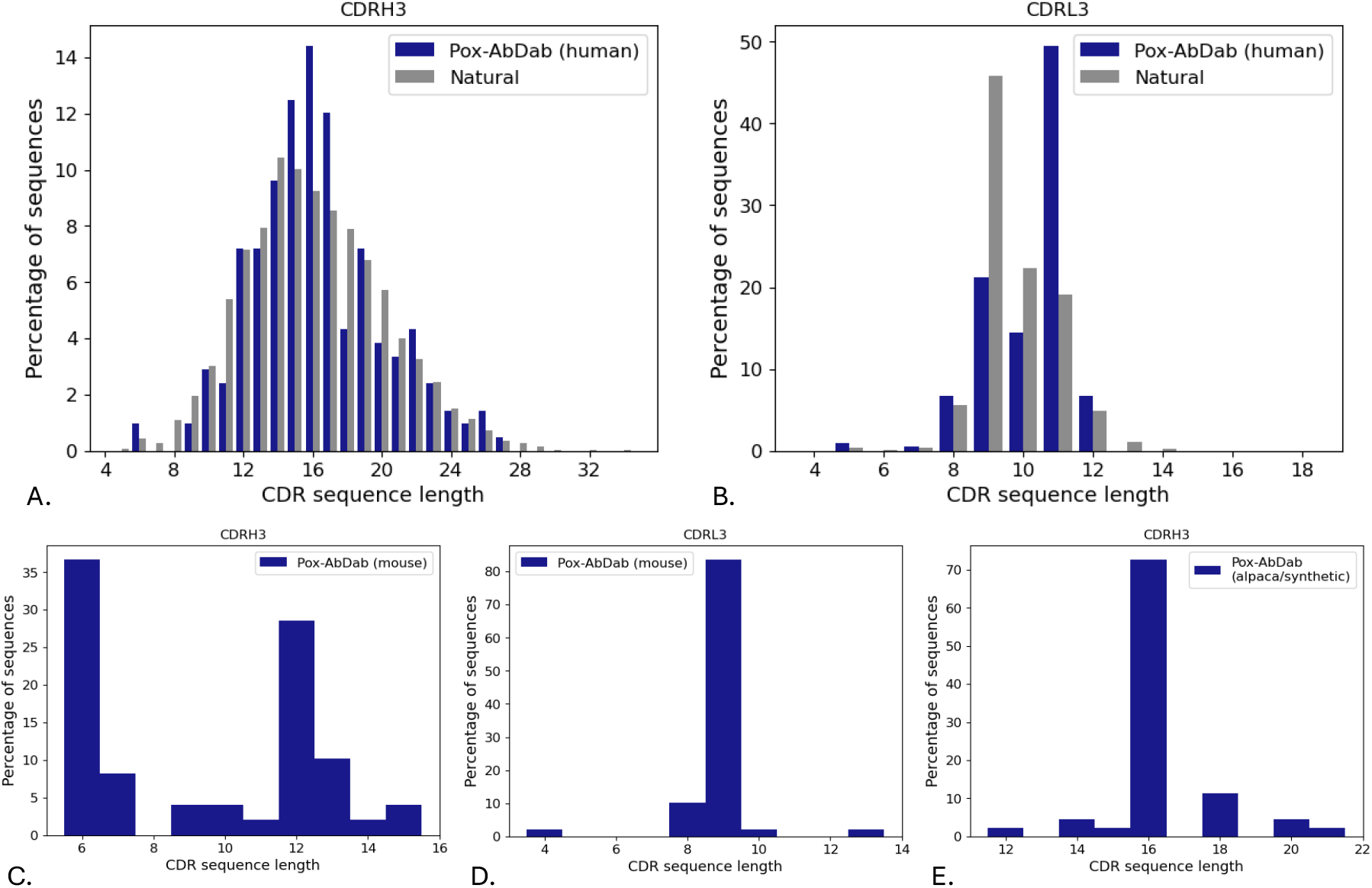
Bar charts showing (A) the CDRH3 and (B) the CDRL3 distributions of all the human antibodies in Pox-AbDab (including those that only have clonotype information) relative to a sample of the baseline natural repertoire from the Observed Antibody Space (OAS) database (38, 39); (C) the CDRH3 and (D) the CDRL3 distributions of all the murine antibodies in Pox-AbDab (including those that only have clonotype information); (E) the CDRH3 distribution of all single-domain antibodies in Pox-AbDab. Comparison against the natural human repertoire is based on 5000 randomly samples natural antibody sequence from the Jaffe et al., 2022 (53) as stored in OAS.

**Figure S6.**
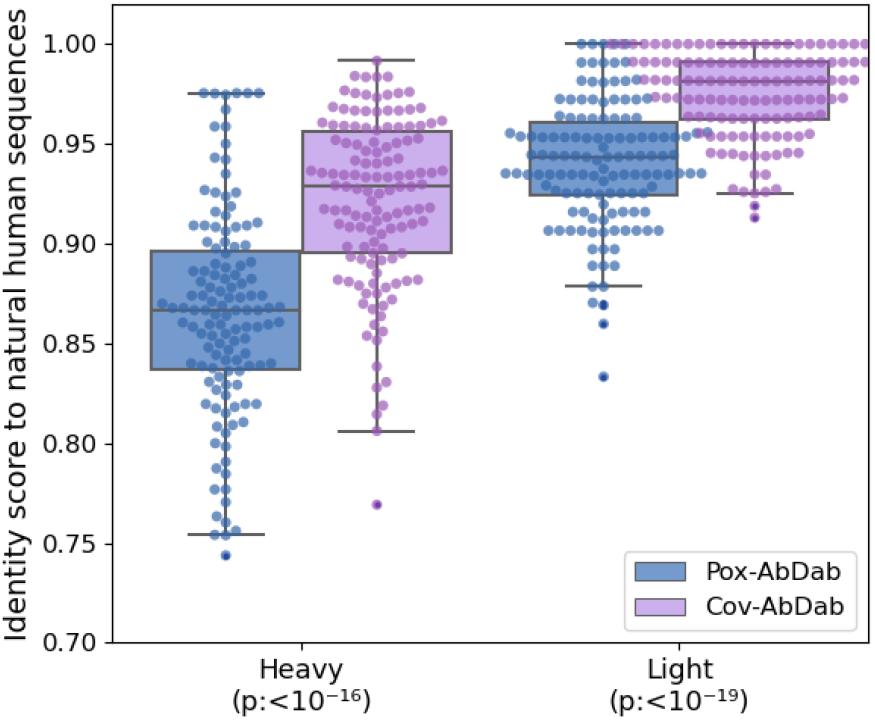
Antibody heavy and light sequence identity score to a set of human antibodies from naive B-cells. The 130 full-length human antibody sequences in Pox-AbDab (blue) and 130 randomly selected human antibody sequences from CoV-AbDab (purple) were compared against all human B cell receptor sequences from OAS using the “OAS-aligned” dataset provided with KA-Search (1.9Bn heavy chains from 69 studies, 350.5M light chains from 38 studies). The highest sequence identity observed was recorded. P-values of the Mann–Whitney U test are indicated on the x-axis.

**Table S1.** VACV proteins with known roles in orthopoxvirus function alongside the data availability for antibodies engaging each protein in Pox-AbDab. All function annotations derive from Moss 2011 (18), except where otherwise indicated.

| Protein in VACV Nomenclature (Strain if not Copenhagen) | Location | Function | Surface protein | Number of targeting antibody sequences | Number of targeting antibody crystal structures |
| --- | --- | --- | --- | --- | --- |
| A10 | MV | Assembly (58) | No | 1 |  |
| A13 | MV | Assembly (59) | Yes |  |  |
| A14 | MV | Morphogenesis (60) | Yes | 5 |  |
| A16 | MV | Entry/Fusion | Yes | 4 |  |
| A17 | MV | Assembly, Fusion | Yes |  |  |
| A21 | MV | Entry/Fusion | Yes | 1 |  |
| WR148 (WR) | MV | Fusion Suppressor | No | 7 |  |
| A26 | MV | Attachment | Yes |  |  |
| A27 | MV | Attachment | Yes | 27 | 2 |
| A28 | MV | Entry/Fusion | Yes | 1 |  |
| A33 | EEV | Spread | Yes | 33 | 3 |
| A34 | EEV | Spread, Attachment (61) | Yes |  |  |
| A56 | EEV | Fusion (62), Immune Evasion (63) | Yes |  |  |
| B5 | EEV | Spread, Morphogenesis | Yes | 27 | 1 |
| C3 | Secreted | Immune evasion | No | 11 |  |
| D8 | MV | Attachment | Yes | 32 | 5 |
| F5 | EEV | Plaque morphology (64) | Yes |  |  |
| F9 | MV | Entry/Fusion | Yes | 1 |  |
| F13 | EEV | Spread (65), Morphogenesis (65) | Yes | 1 |  |
| G3 | MV | Entry/Fusion | Yes |  |  |
| G9 | MV | Entry/Fusion | Yes |  |  |
| H2 | MV | Entry/Fusion | Yes |  |  |
| H3 | MV | Assembly, Attachment | Yes | 39 |  |
| H5 | MV | Morphogenesis (66) | No | 2 |  |
| I1 | MV | Assembly (67) | No | 5 |  |
| J5 | MV | Entry/Fusion | Yes |  |  |
| L1 | MV | Entry/Fusion | Yes | 15 | 2 |
| L5 | MV | Entry/Fusion | Yes |  |  |
| O3 | MV | Entry/Fusion | Yes |  |  |

## Footnotes

1 https://worldhealthorg.shinyapps.io/mpx_global/

2 https://mariadb.com/

3 https://sqlmodel.tiangolo.com/

4 https://spec.openapis.org/oas/v3.1.0.html

5 https://fastapi.tiangolo.com/

